# Surface Expansion Regionalization of the Hippocampus in Early Brain Development

**DOI:** 10.1101/2025.02.22.639699

**Authors:** Ya Wang, Liangjun Chen, Zhengwang Wu, Sheng-Che Hung, J. Keith Smith, Li Wang, Li Tengfei, Weili Lin, Gang Li

## Abstract

The hippocampal formation is implicated in a myriad of crucial functions, particularly centered around memory and emotion, with distinct subdivisions fulfilling specific roles. However, there is no consensus on the spatial organization of these subdivisions, given that the functional connectivity and gene expression-based parcellation along its longitudinal axis differs from the histology-based parcellation along its medial-lateral axis. The dynamic nonuniform surface expansion of the hippocampus during early development reflects the underlying changes of microstructure and functional connectivity, providing important clues on hippocampal subdivisions. Moreover, the thin and convoluted properties bring out the hippocampal maturity largely in the form of expanding surface area. We thus unprecedentedly explore the development-based surface area regionalization and patterns of the hippocampus by leveraging 513 high-quality longitudinal MRI scans during the first two postnatal years. Our findings imply two discrete hippocampal developmental patterns, featuring one pattern of subdivisions along the anterior-posterior axis (head, regions 1 and 5; body, regions 2, 4, 6, and 7; tail, region 3) and the other one along the medial-lateral axis (subiculum, regions 4, 5, and 6; CA fields, regions 1, 2, and 7). Most of the resulting 7 subdivisions exhibit region-specific and nonlinear spatiotemporal surface area expansion patterns with an initial high growth, followed by a transition to low increase. Each subregion displays bilaterally symmetric pattern. The medial portion of the hippocampal head experiences the most rapid surface area expansion. These results provide important references for exploring the fine-grained organization and development of the hippocampus and its intricate cognitions.

## Introduction

The hippocampal formation involves intricate cognitions, such as regulating response inhibition, episodic memory, and spatial navigation (1, 2). Its aberrations have been found in a range of brain disorders (3), such as Alzheimer’s disease, schizophrenia, and temporal lobe epilepsy. There is also evidence suggesting abnormal hippocampal volumes are related to the impairment of social interaction in autism spectrum disorder (ASD) (4, 5). The distinct disease-associated symptoms may stem from the abnormality of different hippocampal subregions relating to specific functions. Therefore, it is of great significance to thoroughly explore the regionalization of normative hippocampal formation and its intricate cognitions.

Traditional histological research emphasizes hippocampal subfields along the medial-lateral axis, encompassing the Cornu Ammonis (CA1-4) subfields, the dentate gyrus, and the subiculum, on the basis of differences in cytoarchitectonic profiles and distinct intrinsic connections among these subfields (6-8). Recently, more attention is being directed towards the internal organization of the hippocampal formation from multiple perspectives (8), given that the disparate hippocampal subregions are under consideration for their differential involvement in cognitive processes (9-12). In contrast to the traditional histology-based hippocampal parcellation, the ensuing view derived from functional profiles and gene expression studies in rodents and nonhuman primates (13-15), and neuroimaging studies in humans in terms of hypothesis-driven activation and connectivity and its relationship to behavior have inspired the proposal of multiple functional domains along the hippocampal longitudinal axis (8, 11, 16, 17), such as a tripartite subdivision (head, body, and tail). However, a paucity of studies delves into hippocampal internal topological organization in terms of early dynamic development, which is essentially related to the underlying changes of both microstructure and functional connectivity, thus providing important clues on hippocampal subdivisions. As the hippocampus is a thin and folded sheet, surface area expansion is well suitable for characterization of its dynamic, complex, spatiotemporally heterogeneous early development.

However, our knowledge of early hippocampal development-based regionalization (the spatial layout of developmentally distinct hippocampal subdivisions) in terms of surface area expansion is still scarce, especially during the first two postnatal years featuring the most dynamic, nonuniform brain development. This primarily arises from challenges associated with the acquisition and processing of infant MR images, which exhibit notably low tissue contrast and dynamic imaging appearance.

Therefore, in this work, we unprecedentedly studied the developmental regionalization of the hippocampal formation in terms of early surface area expansion, by leveraging 513 high-resolution longitudinal MRI scans from 231 subjects densely covering the first two postnatal years. With the aim of capturing the hippocampal configuration, a data-driven non-negative matrix factorization (NMF) (18, 19) method was employed to decompose the hippocampal formation into a set of distinct subregions based on the vertex-wise surface area expansion, where each subregion is formed by a set of co-developing vertices. A spectral clustering (20, 21) was further employed to validate the above parcellation. Accordingly, we charted the region-specific spatiotemporal developmental trajectory of each of our discovered hippocampal subregions and conducted the hierarchical clustering to scrutinize the relationship among these subregions.

## Results

In this study, we first computed the hippocampal vertex-wise surface area based on 513 longitudinal scans from 231 subjects within the first two postnatal years, and then utilized NMF and spectral clustering methods to uncover the hippocampal developmental regionalization map, i.e., the spatial layout of a set of developmentally distinct subregions, based on the local surface area expansion. Finally, we charted and characterized the developmental trajectories of these hippocampal subregions.

### Developmental Regionalization of Hippocampal Surface Area

**Fig. 1** showed the optimal number of hippocampal subregions separately for left and right hippocampi using the NMF method, as determined by local high silhouette coefficients and relatively low reconstruction errors. We observed that the silhouette coefficient reached a local maximum at *r* = 5, 7, and 9 in the left hippocampus (**Fig. 1A**) and the reconstruction error reached a low value and plateau starting from *r* = 7 to 9 (**Fig. 1B**). For the right hippocampus, we observed a high local maximum of silhouette coefficient at *r* = 7 (**Fig. 1C**), and the reconstruction error also exhibited a relatively low value and plateau from *r* = 6 to 8 (**Fig. 1D**). Therefore, we mainly focused on the results at *r* = 7 to represent the hippocampal developmental regionalization. Meanwhile, we also provided the hippocampal partition results when *r* = 2 and 5 to scrutinize the coarse regionalization.

**Fig. 1.**
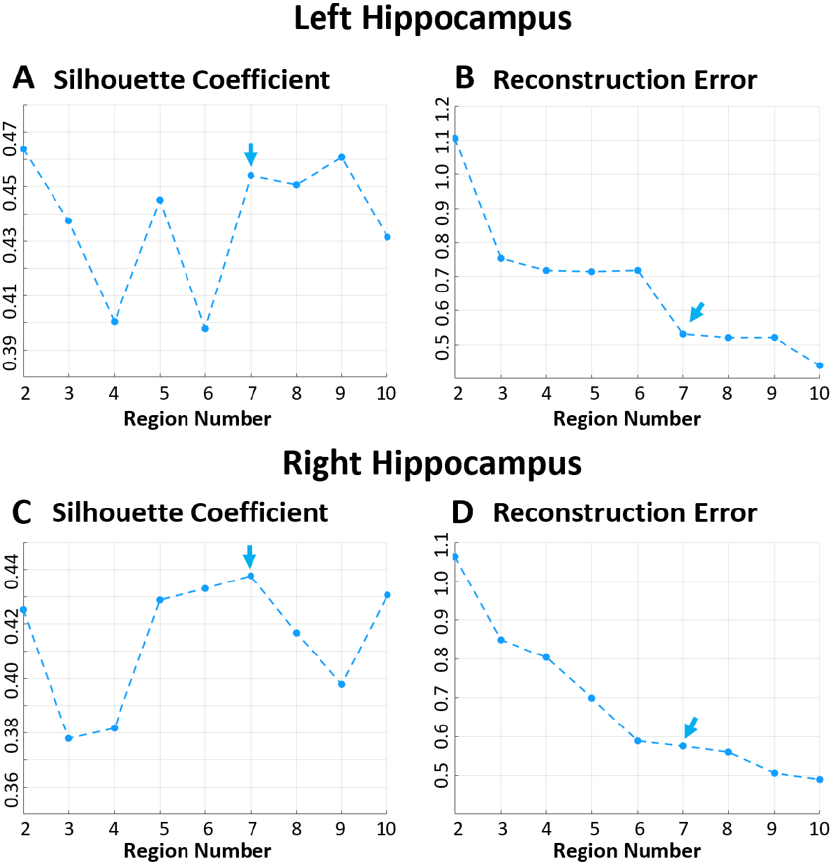
Determine the optimal subregion number *r* in bilateral hippocampi. (**A**) and (**B**) correspond to the average silhouette coefficients and reconstruction errors across varying subregion numbers in left hippocampus. (**C**) and (**D**) pertain to the average silhouette coefficients and reconstruction errors in right hippocampus. The arrows denote the appropriate subregion number with local relatively high silhouette coefficients and low reconstruction errors.

The discovered subregions for *r* = (2, 5, 7) were illustrated in **Fig. 2**. The subregions with *r* = 2 (**Fig. 2A**) within the left and right hippocampi are relatively symmetric, with one subregion predominantly corresponding to the head and the other one encompassing the hippocampal body and tail. When *r* equals to 5 (**Fig. 2B**), despite the lack of symmetry in subregions between the left and right hippocampi, they both exhibit a triple partition. Specifically, within the left hippocampus, the head comprises region 1, the body consists of regions 2, 4, and 5, and the tail encompasses region 3. In the right hippocampus, the head comprises region 1 and 4, the body includes regions 2 and 5, and the tail contains region 3. Nevertheless, when increasing *r* to 7 (**Fig. 2C**), the hippocampal subregions exhibit more bilaterally symmetric patterns than that with fewer subregions. Specifically, two distinct developmental patterns of hippocampal surface area are discerned, with one pattern adhering to the tripartite subdivision, encompassing hippocampal head (regions 1 and 5), body (regions 2, 4, 6, and 7), and tail (region 3). The other pattern follows a medial-lateral partition, with the medial portion consisting of regions 4, 5, and 6, while the lateral portion comprising regions 1, 2, and 7. Of note, there is no specific medial-lateral division for region 3. The medial regions primarily map to the subiculum, while the lateral regions predominantly correspond to the hippocampal CA1-4 and the dentate gyrus. The spatial positions of the hippocampus relative to the brain was shown in **Fig. 2D**.

**Fig. 2.**
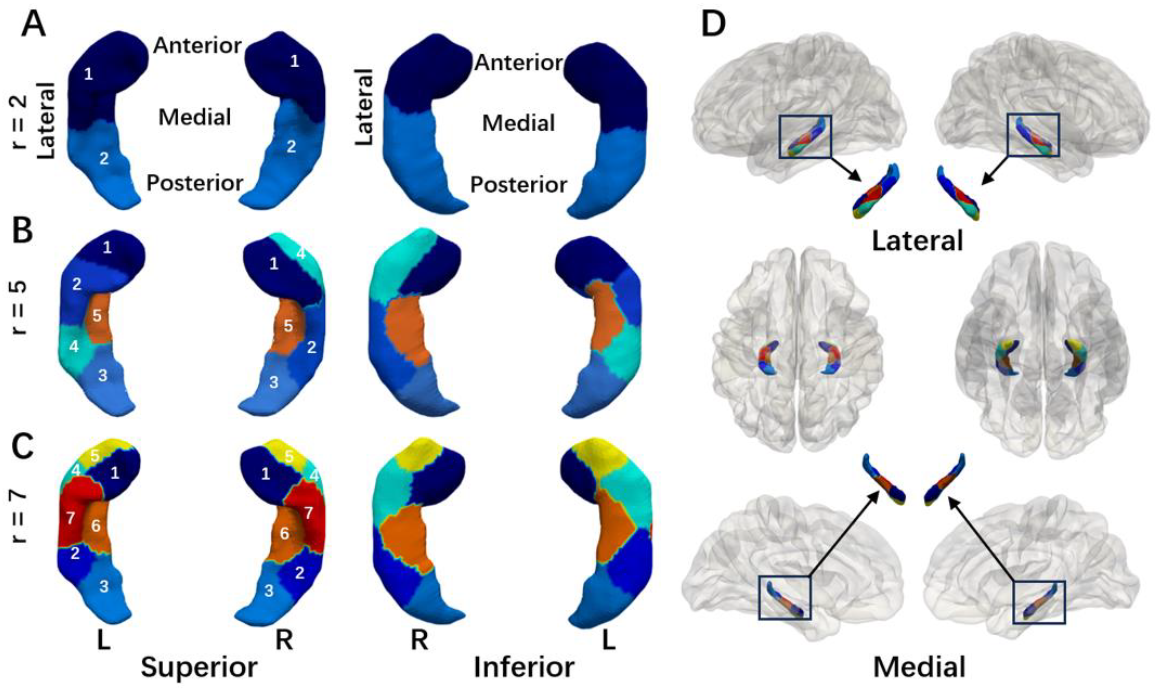
The developmental regionalization maps of surface area. Panels (**A**), (**B**), and (**C**) showcase the hippocampal parcellation maps with different region numbers (*r* = 2, 5, and 7) in bilateral hippocampi. (**D**) The spatial position of hippocampus relative to the brain. The coloring in **D** corresponds to **C** (r = 7).

The optimal numbers of hippocampal subregions for left and right hippocampi using the spectral clustering method were shown in **Fig. S1**. In the left hippocampus, when *r* equals 5 and 7, the silhouette coefficient reached its local maximum. We thus opted for *r* = 7, in alignment with the above results from NMF, to portray the hippocampal regionalization, despite that it may not be the optimal subregion number in the right hippocampus according to the silhouette coefficient. The spectral clustering-based developmental regionalization of surface area in left and right hippocampi is also relatively symmetric and exhibits two kinds of developmental patterns (**Fig. S2**), aligning with the results obtained by NMF.

**Fig. 3A** displayed the dendrogram of the hierarchical relationship among the hippocampal subregions. Accordingly, the 7 hippocampal subregions were categorized into 3 groups, i.e., hippocampal head (regions 1 and 5), body (regions 2, 4, 6, and 7), and tail (region 3) (**Fig. 3B**).

**Fig. 3.**
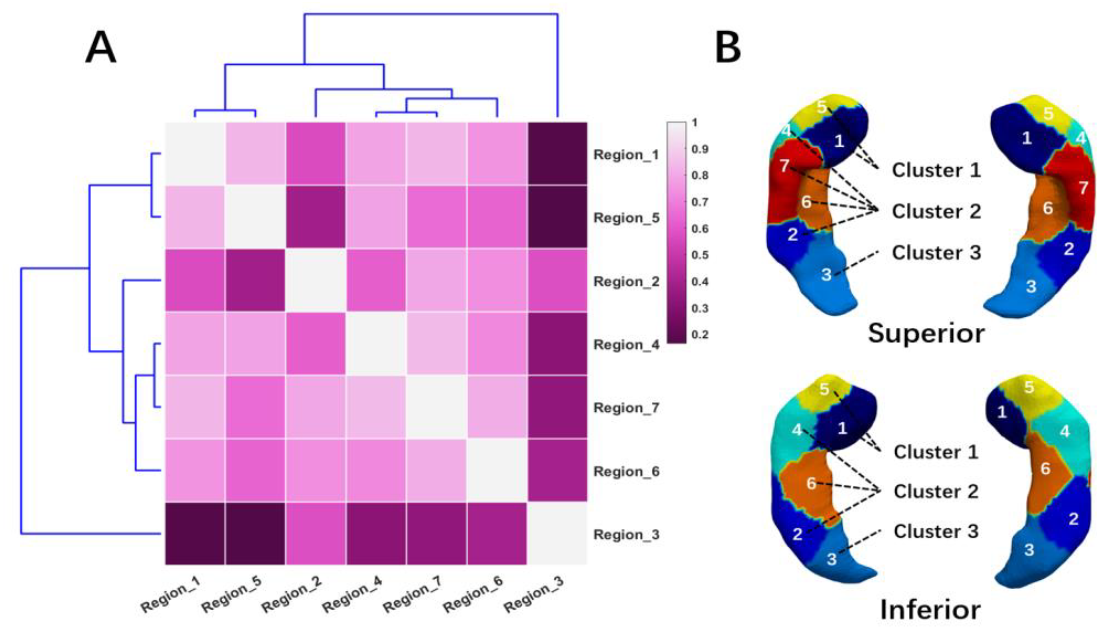
Organization among the hippocampal subregions. The dendrogram of the hierarchical relationship (**A**) among the hippocampal subregions using the Ward D2 method and the visualization upon grouping these regions into three distinct classes (**B**).

### Developmental Patterns of Discovered Hippocampal Subregions

The population-level longitudinal developmental trajectory of the hippocampal surface area in each discovered subregion fitted by GAMM is shown in **Fig. 4** (**A**, left; **B**, right). Meanwhile, the normalized surface area compared to the surface area at birth and the monthly expansion velocity of the surface area in each hippocampal subregion at 1, 2, 3, 4, 5, 6, 9, 12, 18, and 24 months are depicted in **Fig. 5** (**A**) and (**B**), respectively. As per our findings, the hippocampal surface area exhibits region-specific and bilaterally relatively symmetric developmental patterns within the first two postnatal years. Except for regions 2 and 3, which exhibit relatively low and nearly linear surface area expansion, the absolute surface areas of most subregions in both hippocampi adhere to a nonlinear growth pattern marked by an initial phase of high expansion, followed by a subsequent transition to low expansion. No peaks arrive during the whole age ranges examined. Among these subregions, region 1 (the medial portion of the hippocampal head) experiences the most rapid surface area expansion. Furthermore, the hippocampal subregions showed remarkable sex differences, with males having larger absolute surface area than females. The detailed age ranges with significant sex differences were shown in **Table S1**.

**Fig. 4.**
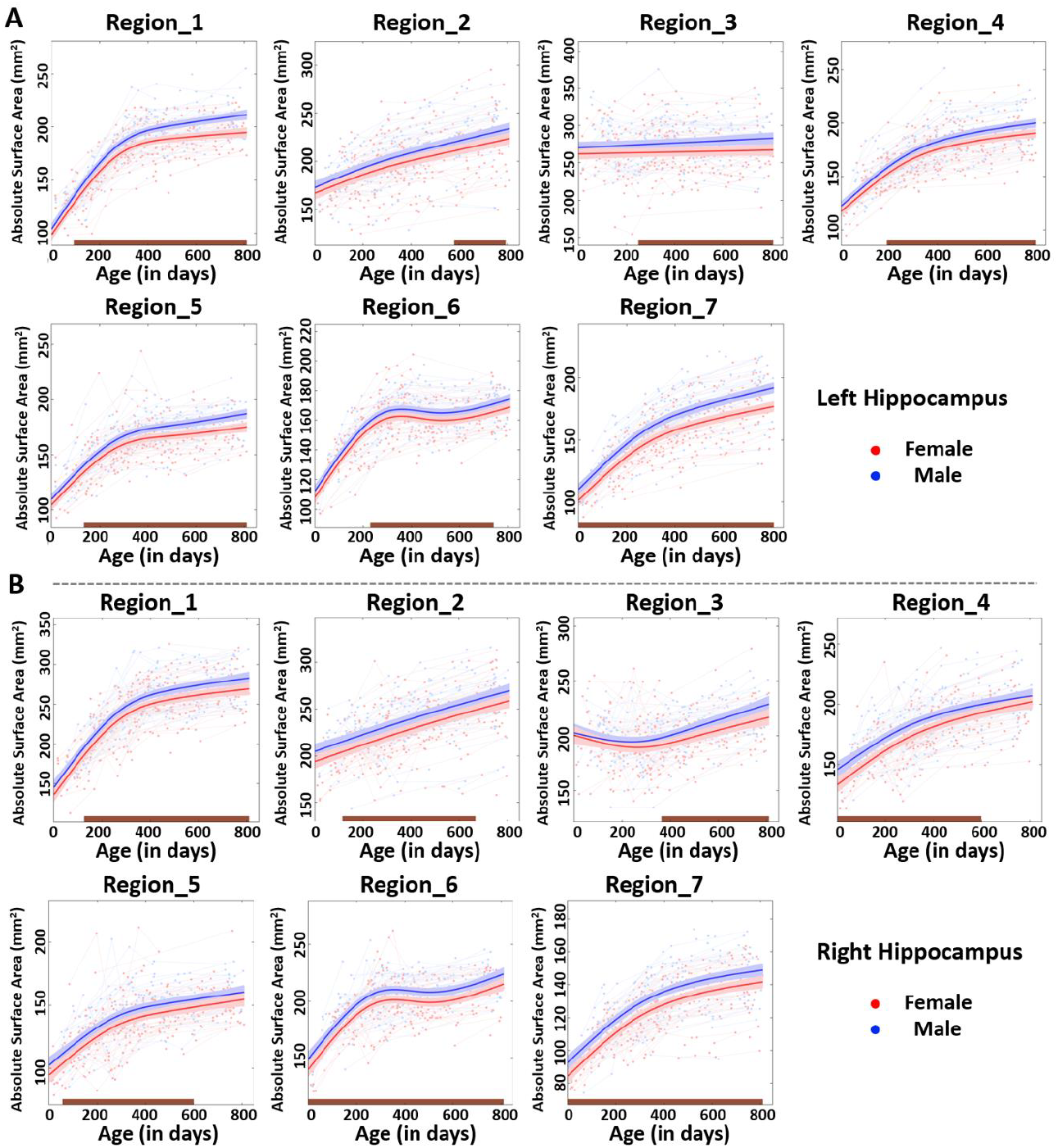
The longitudinal developmental trajectories of hippocampal *absolute* surface area. (**A**) and (**B**) separately depict the surface area developmental pattern of each subregion in left and right hippocampi. The solid red and blue curves illustrate the population-fitted trajectories of each hippocampal subregion for females and males, respectively. The bule and red faint lines represent the developmental trajectories of each subject with multiple scans for males and females, respectively. The shaded ribbons represent 95% confidence intervals of the fitted curves. The brown horizontal bars on the X-axis mark the age intervals with significant sex differences.

**Fig. 5.**
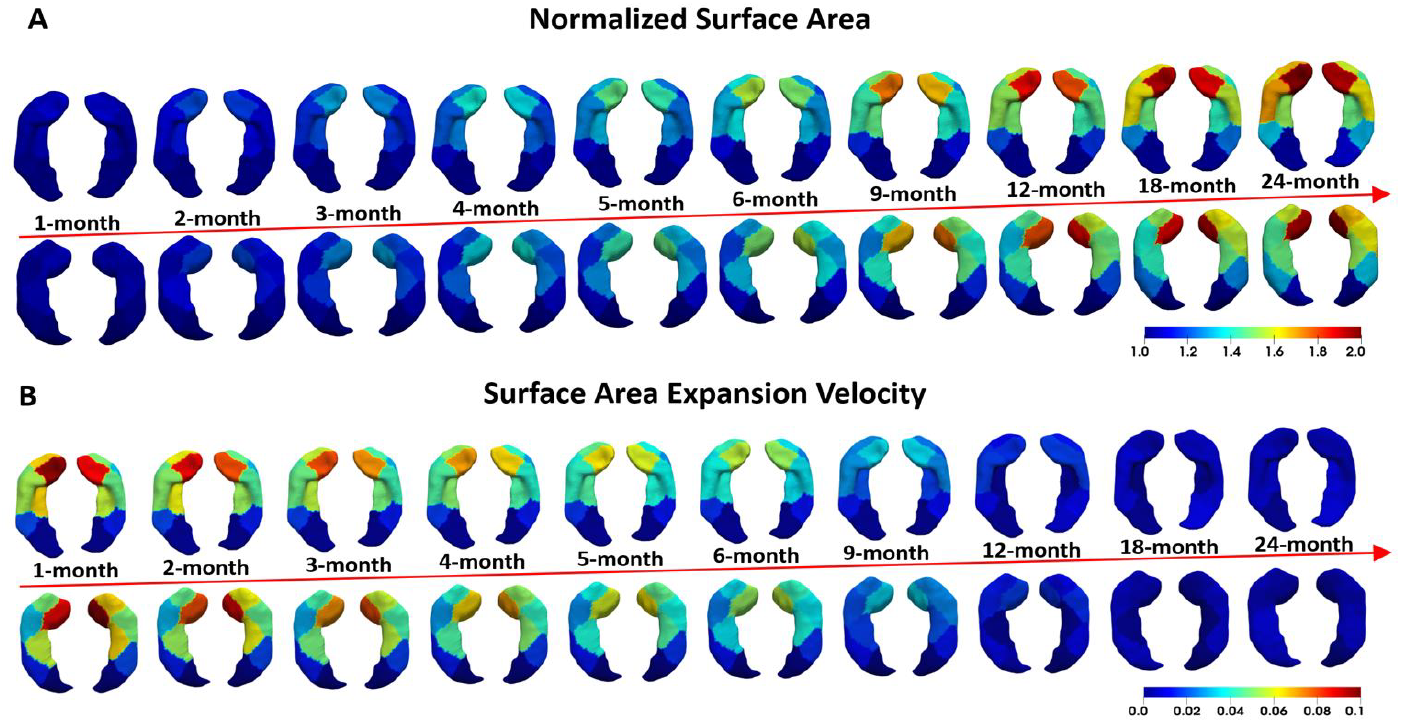
The normalized surface area and the surface area expansion velocity. (**A**) and (**B**) respectively showcase the normalized surface area relative to the surface area at birth and the monthly expansion velocity of the surface area in each hippocampal subregion at time points 1, 2, 3, 4, 5, 6, 9, 12, 18, and 24 months.

**Fig. 6** illustrated the population-level longitudinal developmental trajectories of hippocampal relative surface area in each subregion of both left (**Fig. 6A**) and right (**Fig. 6B**) hippocampi, respectively. As observed, regions 1 and 5 (head) showed more rapid surface area expansion relative to the ipsilateral hippocampal formation during the first postnatal year and then transitioned into a slow expansion phase within the second postnatal year. In contrast, regions 4 and 7 (body) exhibit a prolonged phase of rapid surface area expansion relative to the ipsilateral hippocampal formation. However, the surface areas of regions 2 and 3 (tail) exhibited slower expansions than that of the ipsilateral hippocampal formation first and then this relative slowness gradually reduced. Differing from other subregions, region 6 displayed a more rapid surface area expansion followed by a slower expansion than the ipsilateral hippocampal formation. Of note, most subregions lack significant sex differences except for left regions 6 and 7. The relative surface area of region 6 exhibited larger expansion in females than males (age ranges: 326-810 days), and region 7 showed larger surface area expansion in males than females (age ranges: 0-505 days).

**Fig. 6.**
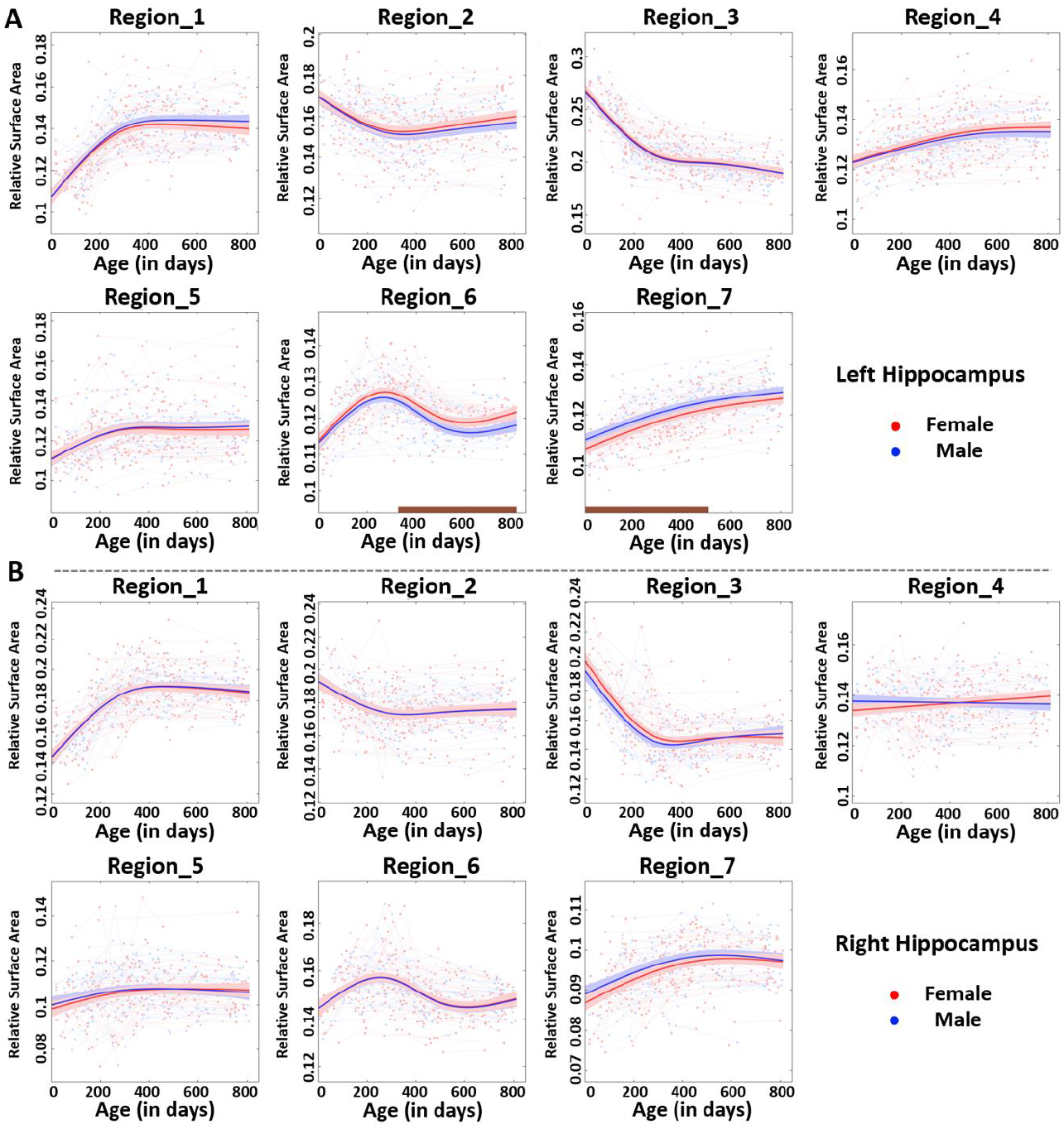
The longitudinal developmental trajectories of hippocampal *relative* surface area. (**A**) and (**B**) separately delineate the surface area developmental pattern of each subregion in left and right hippocampi. The solid red and blue curves depict the population-fitted trajectories in each hippocampal subregion for females and males, respectively. The blue and red faint lines represent individual subject-specific surface area developmental trajectories, for males and females, respectively. The shaded ribbons represent 95% confidence intervals of the fitted curves. The brown horizontal bars on the X-axis delineate the age intervals with notable sex differences.

## Discussion

This is the first study to explore the hippocampal regionalization from a longitudinal developmental perspective, employing large-scale data from subjects within the first two postnatal years. By discovering clusters of vertices that are co-developing across ages and subjects using data-driven clustering methods, our findings imply that the developing hippocampal formation displays two discrete organizational patterns, featuring one pattern of subdivisions along the anterior-posterior (AP) axis and the other one along the medial-lateral (ML) axis. In the anterior-posterior pattern, the hippocampus is mainly partitioned into the head (regions 1 and 5), body (regions 2, 4, 6, and 7), and tail (region 3). Along the medial-lateral and ventrodorsal axes, the hippocampus is parcellated into subdivisions corresponding to CA1-4 (regions 1, 2, and 7) and subiculum (regions 4, 5, and 6). Each portion of these resulting 7 subdivisions show a relatively symmetric pattern of surface area expansion in bilateral hippocampi during infancy. The further hierarchical clustering grouped these 7 hippocampal subregions into head, body, and tail portions. In conjunction with the aforementioned outcomes, we also unveil that each subregion presents its own spatiotemporally distinct surface area expansion. This work proposes an important hippocampal parcellation method derived from surface area expansion during early postnatal years. Charting the developmental trajectories of surface area in each normal hippocampal subregion provides a valuable reference for morphological abnormality occurring during early neurodevelopmental disorders.

A plethora of early studies pertaining to the hippocampal organization indicate the hippocampal formation following one single pattern, either along the mediolateral axis or the long axis according to the publications (7, 8, 22, 23). However, increasing evidence supports hippocampal organization adheres to both patterns and even continuous gradients within hippocampal internal structure (24-27). Our results suggest the hippocampal regionalization patterns from developmental perspective in infants follow two dimensions. This may be attributed to the discrepancies of structure, function, and connectivity orchestrated by hippocampal subregions. As to along the medial-lateral axis, apart from varying cellular architectures along the medial-lateral axis (28, 29), the established crucial neural circuits by these subfields are also the reflection of distinct functions among each subfield. For instance, the classic tri-synaptic pathway, comprised of the dentate gyrus, CA3, and CA1, receives input from the entorhinal cortex while projecting efferent signals mainly via the subiculum (28-30). Each subfield along the medial-lateral axis exhibits distinct functions. Moreover, recent functional connectivity research in adults using MRI has also revealed that different subfields along the medial-lateral axis exhibit their unique patterns of functional connectivity (10, 11) as well as different portions along the longitudinal axis (11, 17, 31, 32), which is in line with our findings in infants. The partition patterns along the anterior-posterior axis may be related to the specific cognitions. A recent study using multimodal connectivity-based parcellation and behavioral profiling revealed an emotion-cognition gradient as well as a self-world-centric gradient along the longitudinal axis in adults (25). The above conjectures need to be further validated in infants.

After fitting the developmental trajectory of the surface area in each hippocampal subregion, we find that most subregions (except for regions 2 and 3) exhibit remarkable absolute surface area expansion followed by a sluggish increase, which aligns with the developmental trajectory of hippocampal volume within the first two postnatal years (33-35). One significant factor for the rapid development might be its involvement in the establishment of structural connectivity, such as the monosynaptic pathway, during the early postnatal years (36). Parallel to histological development, behavioral progress suggests that neither pathway (monosynaptic and tri-synaptic pathway) is full established during infancy and the first basic forms of episodic memory was provided by the monosynaptic circuit during the second postnatal year (36). Moreover, our results reveal that the hippocampal head undergoes the most rapid growth followed by the body and the tail shows the slowest growth during the first two postnatal years. Hippocampal shape analysis studies also suggest a different expansion pattern within the hippocampus with the nonlinear, rapid expansion mainly happened in the hippocampal head during childhood (37, 38). This distinctive expansion pattern may be related to the detailed functional formation along the anterior-posterior axis (25, 37) and requires further confirmation.

The absolute surface areas of most subregions exhibit significant sex differences during different age ranges, with males manifesting larger surface area than females, while nearly no significant sex differences appear in terms of relative surface area expansion (except for left regions 6 and 7). Previous studies on hippocampal volumes also suggest a larger absolute hippocampal volume in males than females in infancy and adults (34, 39), while the assertion regarding the sex difference becomes inconsistent when accounting for overall brain volume (40, 41). Considering relatively limited data in each month especially the first several postnatal months, further studies with multiple datasets with substantial amount of data can confirm our results.

In conclusion, this work unprecedentedly explored hippocampal regionalization from a developmental perspective based on surface area expansion in infants during the first two postnatal years. Our results suggest that the early developing hippocampus contains multiple developmentally distinct subregions, which display two discrete organizational patterns along the anterior-posterior axis and the medial-lateral axis, respectively. Each subregion exhibits its own spatiotemporally distinct surface area expansion patterns, with the medial portion of the hippocampal head undergoing the highest growth, while the tail showing the lowest expansion. The differential surface area developmental patterns across subregions may partly attribute to the establishment of monosynaptic circuit for specific functions during the first two postnatal years. Furthermore, we also reveal relatively symmetric developmental patterns in bilateral hippocampal subregions and more rapid surface area expansion in males than females. Overall, this work provides valuable reference for exploring finer-grained hippocampal structural and functional organizations and their abnormalities during early postnatal years.

## Limitations

Certain limitations need to be noted in this study. First, there are few scans within the first month after birth, which may impact the accuracy of fitting the longitudinal trajectories and the age ranges with significant sex differences of hippocampal subregions. Second, despite we employed two clustering methods for discovering hippocampal developmental regionalization, other datasets within the same age group should be employed to further validate our results. Third, while we have delineated the dynamic longitudinal developmental trajectories within the first two postnatal years, it would be beneficial to extend these development trajectories to cover a broader age range, including prenatal, childhood, adulthood, to full grasp the developmental trajectories and patterns of the hippocampus throughout the life span in our future work.

## Methods

### Human Subjects

The high-quality MR images from healthy subjects utilized in this work were from the UNC/UMN Baby Connectome Project (BCP) dataset (42), which recruited typically developing infants, toddlers, and preschool-aged children, ranging from term birth to 5 years of age. Approval for all procedures was obtained from the Institutional Review Boards of the University of North Carolina at Chapel Hill and the University of Minnesota. Meanwhile, informed consents were signed by the parents of all participants before enrolled. Participants were included under the following criteria: 1) born between 37 and 42 weeks of gestational age, 2) experienced no major pregnancy or delivery complications, 3) birth weight fitting for the gestation age. For an extensive list of inclusion and exclusion criteria, kindly refer to Howell et al. (42). Totally, 702 scans were acquired before starting this study. Comprehensive participant characteristics can be found in **Fig. S3**, while a monthly breakdown of scan numbers is available in the reference (34).

### Image Acquisition

All the MR images were acquired on 3T Siemens MRI scanners using a 32-channel head coil at the Biomedical Research Imaging Center (BRIC) at the University of North Carolina at Chapel Hill and the Magnetic Resonance Research Center at the University of Minnesota. The same scanning sequences, imaging protocols, and quality control criteria were used to mitigate the discrepancies between the two sites. The scans were conducted after children fell asleep naturally, with earplugs in place for ear protection and foam pieces positioned to provide head support while preventing motion artifact due to respiration-related motion. The entire scanning process is monitored by a well-trained expert. The T1-weighted and T2-weighted (T1w and T2w) images were obtained via MPRAGE and variable flip angle turbo spin-echo sequence, with the subsequent parameters: matrix = 320×320, field of view (FOV) = 256×256, resolution = 0.8×0.8×0.8 mm^3^ and 208 sagittal slices. The TR/TE (repetition time/echo time) parameters for T1w and T2w images were 2400/2.24 ms and 3200/564 ms, respectively. All images underwent visual inspection-based quality control by one anatomical expert to exclude those with motion artifacts, inadequate coverage, and/or ghosting. As a result, 564 scans encompassing both T1w and T2w images were remained. For more details, please refer to the reference (34).

### Image Processing

All the scans were processed with the infant brain extraction and analysis toolbox (iBEAT V2.0) (http://www.ibeat.cloud/) (43). Specifically, the N3 method (44) was used to correct the intensity inhomogeneity. Following this, a trained densely connected U-net model (45) was utilized to perform the skull stripping. Then, we employed the FLIRT (46) in FSL to linearly align each T2w image onto its corresponding T1w image. After these processing steps, a total of 513 scans survived for further analysis.

To map the surface area regionalization of the hippocampal formation, we utilized our 4D infant brain volumetric atlas (including age-specific atlases at 0, 1, 2, 3, 4, 5, 6, 7, 8, 9, 10, 11, 12, 15, 18, 21, and 24 months of age) (47), which provide manually delineated hippocampal labels (48). When building the 4D atlases, we obtained the deformations between neighboring age-specific atlases and between each age-specific atlas and the age-matched individual scans (34, 47, 49). To obtain the vertex-to-vertex correspondences of the hippocampal surfaces across subjects and ages, we reconstructed the hippocampal surface mesh representation in 0-month atlas and then warped it to each individual scan at each age by combining the corresponding deformations between age-specific atlases and those between each age-specific atlas and the age-matched scans. For example, for a 2-month individual scan, we concatenated the deformation fields in the following order: 0-month atlas to 1-month atlas, 1-month atlas to 2-month atlas, 2-month atlas to 2-month individual scan. Finally, the local surface area of each vertex was calculated on each individual hippocampal surface and then smoothed. Subsequently, a data matrix was formed for both left and right hippocampal formations with each column indicating the local surface areas of all vertices of an individual hippocampal surface and each row corresponding to the local surface areas of all scans at the same vertex.

### Discovering Distinct Subregions with Non-negative Matrix Factorization

To reveal the spatiotemporal regionalization of the hippocampal surface during infancy, we adopted a data-driven, hypothesis-free method, non-negative matrix factorization (NMF) (18), to partition the hippocampal surface into a set of developmentally distinct regions by clustering the co-developing hippocampal vertices into same regions. This NMF method has been successfully used for the parcellation of brain networks and cortical surface (50-52). The merit of NMF is its adherence to non-negative entries and only additive combinations, allowing NMF learns a part-based representation across all subjects and ages, thereby facilitating the interpretation of discovered hippocampal subregions.

Specifically, in NMF, the non-negative data matrix *V* is a linear combination of the columns of a non-negative base/component matrix *W* weighted by the rows of a coefficient matrix *H*. The equation is abbreviated as *V* ≈ *WH* and NMF can be calculated as min 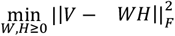. The columns of the matrix *W* are commonly interpreted as representing the components, clusters, or regions in alignment with the research objectives.

In our study, the large non-negative data matrix *V* ∈ *R*^*m*×*n*^ consists of local surface area values for the left and right hippocampal formation separately, where *m* and *n* depict the number of unilateral hippocampal surface vertices and the number of scans. After performing

NMF, the data matrix *V* is decomposed into matrices *Wm*×*r* and *Hr*×*n*, where *r* is generally small (*r* ≪ min{*m, n*}), to achieve dimensionality reduction. The resulting non-negative elements in each column of the matrix *W* intuitively identify a set of vertices undergoing joint development across ages, thereby signifying a distinct region during the developmental regionalization. The corresponding *r* thus denotes the number of hippocampal subregions. As NMF method requires a predefined number of clusters, we set an adequate *r* according to previous hippocampal parcellation.

### Determination of Subregion Numbers

We utilized the silhouette coefficient and reconstruction error jointly to obtain the appropriate hippocampal subregion numbers *r* in NMF.

#### Reconstruction Error

The reconstruction error pertains to the variance between the original data and the data reconstructed by a model. Here, the reconstruction error was calculated as the distance between the original matrix *V* and the matrix reconstructed by identified components and coefficients 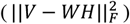, thus a well-suited region number should correspond to a relatively low reconstruction error.

#### Silhouette Coefficient

Two crucial factors are commonly used to evaluate the clustering results, i.e., intraclass dissimilarity and interclass dissimilarity. Intraclass dissimilarity gauges how distinct the objects within the same cluster, whereas interclass dissimilarity quantifies how distinct one cluster is from other clusters. Thus, an optimal clustering solution exhibits low intraclass dissimilarity and high interclass dissimilarity. Here, as in references (20, 53), the silhouette coefficient combining these two factors is adopted to evaluate the NMF clustering results, which is computed as: *sci =* (*bi* − *ai*)/max (*ai, bi*), where *sci* is defined as the silhouette coefficient of the vertex *i, ai* stands for the average dissimilarity between the vertex *i* and all other vertices within the same cluster, and *bi* refers to the minimum average dissimilarity between the vertex *i* and vertices in clusters which the vertex *i* doesn’t belong to. Herein, the dissimilarity between two vertices is calculated by subtracting their Pearson’s correlation of their surface area values across all scans from one. The average silhouette coefficient of all vertices was used, with higher average silhouette coefficients signifying better results.

### Spectral Clustering Analysis

To validate the stability of the discovered developmental regionalization of hippocampal surface area, we utilized another fundamentally different clustering method, spectral clustering (20, 21), to perform further cluster analysis from a different vantage point, by leveraging the correlation between the surface area of each hippocampal vertex and each of 36 cerebral cortical regions from the corresponding scans (51). Specifically, we obtained the surface area value of each cerebral cortical region in both hemispheres parcellated based on the cortical surface area expansion from our previous work (51). After normalization through division by the total surface area of the hippocampus and cerebral cortex within each scan separately, we calculated the Pearson’s correlation of surface area between each hippocampal vertex and each cerebral cortical region and then generated correlation matrices for bilateral hippocampi, which were subsequently utilized to construct the similarity matrices, serving as input to perform the spectral clustering (53).

Here, spectral clustering was applied due to its stronger adaptability to data distribution and much less computational complexity in clustering by mapping the data into an eigen space and can fully utilize the topological relationship among samples (21).

### Statistical Analysis of Developmental Trajectories

After ascertaining the proper number of subregions based on the hippocampal surface area expansion, we employed the non-parametric model, known as generalized additive mixed model (GAMM) (54), to fit the developmental trajectory of hippocampal surface area in each discovered region. This model fits nonlinear and dynamic trajectories better than parametric models in a data-driven way (19). The non-parametric GAMM model is defined as:

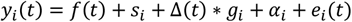

where the two non-parametric functions *f*(*t*) and Δ(*t*) were fitted with the cubic splines; *si* represents the site information; *gi* signifies the sex information (1 for males and 0 for females) of the *i*-th subject; *αi* is the random intercept effect for the *i*-th subject; *ei*(*t*) refers to the random Gaussian noise for the *i*-th subject at time *t* (*i* = 1, 2, …, n and *t* = 0, 1, …, 810 (in days)). For the sex difference, Simpson’s method (55) is employed to calculate the simultaneous confidence interval of the fitted curves between male and female infants at each time point for each hippocampal subregion. The age ranges where the simultaneous confidence interval does not encompass zero are regarded as having significant sex difference. Following that, we calculated the normalized surface area of each subregion based on the corresponding values at day 0, with respect to the differences in sex. Afterwards, we obtained the monthly expansion velocity of the surface area in each hippocampal subregion according to the fitted trajectory. To delve into the relationship among the hippocampal subregions, we utilized the dendrogram to depict the hierarchical organization of the hippocampal subregions as employed in previous studies (53, 56). Specifically, we first computed the Pearson’s correlation among the 7 hippocampal subregions based on the surface area across subjects and scans after averaging the left and right hippocampi. After transforming the correlation matrix into a distance matrix, we computed the hierarchical relationships among the hippocampal subregions.

## Acknowledgements

This work was supported in part by NIH grants (MH116225, MH117943, MH104324, MH123202, and MH116527) and the National Institute on Aging (NIA) of the National Institutes of Health (NIH) under Award Number RF1AG082938 and U01AG079847. This work also utilizes approaches developed by an NIH grant (1U01MH110274) and the efforts of the UNC/UMN Baby Connectome Project Consortium.

## Author contributions

Y.W., L.C., and G.L., investigation, methodology, validation; Y.W., L.C., Z.W., L.W., W.L., and T.L., data curation, formal analysis, software; G.L., funding acquisition, supervision, and conceptualization; Y.W., L.C., S.H., K.S., and G.L., writing, visualization.

## Data availability

All the original BCP data are from publicly available databases (https://nda.nih.gov/edit_collection.html?id=2848). The source data mentioned in the main text and the supplementary materials are available. The 4D infant brain volumetric atlas used in this study is also publicly available (https://www.nitrc.org/projects/uncbcp_4d_atlas/). Other data that support the findings of this study are available on request from the corresponding author [G.L.].

## Code availability

The MR image processing was carried out using iBEAT V2.0 (http://www.ibeat.cloud/). Advanced Normalization Tools (ANTs 2.2.0, https://github.com/ANTsX/ANTs) were utilized for image registration. The NMF, spectral clustering, and curve figures were performed in MATLAB 2023b. The GAMM used in statistical analysis was performed in R 4.1.3 (primarily packages: gam 1.22-2, https://cran.r-project.org/web/packages/gam/index.html). Other codes or software used in this study are available from the corresponding author.

