## Supplementary Materials for "Surface Expansion Regionalization of the Hippocampus in Early Brain Development"

### Surface Expansion Regionalization of the Hippocampal Formation in Early Brain Development

Ya Wang *et al.*

**This PDF file includes:**

Figs. S1 to S3
Tables S1

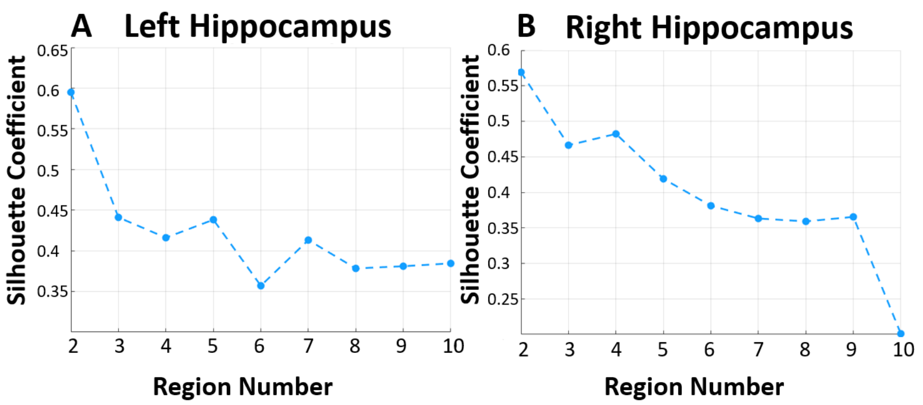


**Fig. S1**. **The silhouette coefficients of hippocampus at different subregion numbers.** (**A**) and (**B**) separately signify the silhouette coefficients of hippocampal surface area within distinct subregions calculated using spectral clustering method.

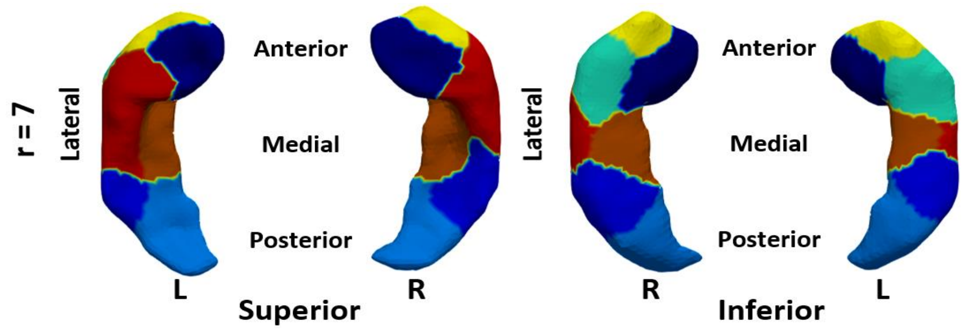


**Fig. S2**. **The developmental regionalization maps of surface area with 7 subregions.** The parcellation map in both left and right hippocampi achieved using spectral clustering method.

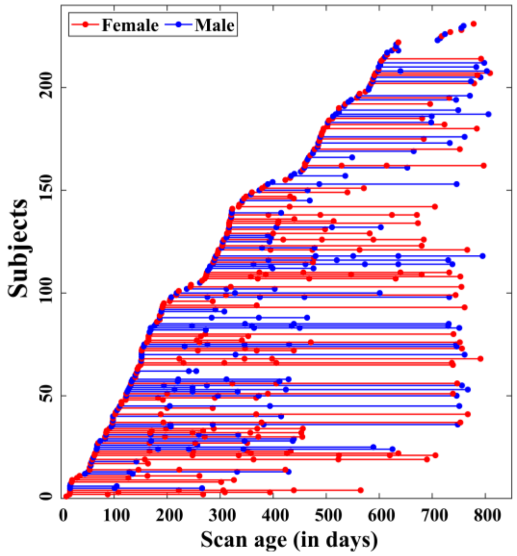


**Fig. S3**. **The longitudinal distribution of scan ages.** Each horizontal line corresponds to one subject and each point denotes one scan at its corresponding scanning age. Blue: male; red: female.

**Table S1. The age ranges with significant sex differences of absolute surface area in both left and right hippocampal subregions.** R, region; LH, left hippocampus; RH, right hippocampus.

| | Age Ranges (days) | | | | | | |
|---|---|---|---|---|---|---|---|
| | **R_1** | **R_2** | **R_3** | **R_4** | **R_5** | **R_6** | **R_7** |
| **LH** | 92-810 | 579-793 | 249-810 | 189-810 | 134-810 | 229-741 | 0-810 |
| **RH** | 125-810 | 114-668 | 363-810 | 0-598 | 55-600 | 0-810 | 0-810 |
